# Fibrous caps in atherosclerosis form by Notch-dependent mechanisms common to arterial media development

**DOI:** 10.1101/2020.09.28.316984

**Authors:** Carlos J. Martos, Julián Albarrán-Juárez, Daniel Morales-Cano, Ainoa Caballero, Donal MacGrogan, José Luis de la Pompa, Laura Carramolino, Jacob F. Bentzon

## Abstract

**Rationale:** The rupture of the fibrous cap in atherosclerotic lesions is the underlying cause of most thrombi leading to heart attack and a frequent cause of stroke. Caps are produced by smooth muscle cells (SMCs) that are recruited to the subendothelial space. We hypothesized that the recruitment mechanisms are likely common to embryonic artery development, which relies prominently on Notch signaling to form the subendothelial layers of medial SMCs.

**Objective:** To analyze the causal roles of the Notch signaling pathway in SMCs for atherogenesis and cap formation.

**Methods and Results:** Notch elements involved in arterial media development were found expressed in regions of fibrous cap in mouse plaques. To assess the causal role of Notch signaling in cap formation, we studied atherosclerosis in mice in which the Notch pathway was inactivated specifically in SMCs by conditional knockout of the essential effector transcription factor RBPJ. No major effects were observed on plaque size, but the presence of cap SMCs was significantly reduced. Lineage tracing revealed that the accumulation of SMC-derived plaque cells in the cap region was unaltered but that Notch-defective cells failed to re-acquire the SMC phenotype in the cap. To analyze whether the accumulation of SMC-derived cells in atherogenesis requires down-regulation of Notch signaling, we studied atherosclerosis in mice with constitutive Notch signaling in SMCs achieved by conditional expression of the Notch intracellular domain. Forced Notch signaling inhibited the ability of medial SMCs to contribute to plaque cells, including both cap SMCs and osteochondrogenic cells, and significantly reduced atherosclerosis development.

**Conclusions:** Sequential loss and gain of Notch signaling is needed to build the cap SMC population. The shared mechanisms with embryonic arterial media assembly suggest that the fibrous cap forms as a *neo-media* that restores the connection between endothelium and stabilizing SMCs, which is transiently disrupted by atherogenesis.

## Introduction

The fibrous cap of atherosclerotic lesions is a key determinant of the clinical course of atherosclerosis. Only lesions with degraded or poorly formed caps (thin-cap fibroatheromas) are at risk of undergoing plaque rupture, the process whereby a tear in the cap exposes necrotic plaque material to the hemostatic system, leading to luminal thrombosis and heart attack or ischemic stroke (1).

Caps are produced by SMCs that are recruited to the subendothelial space during lesion development. Recent work by our group and others has shown that cap SMCs are clonally expanding sheets of cells that spread under the endothelium (2,3). This process restores the intimate relationship between endothelium and SMCs that normally exists throughout the arterial tree but is transiently disrupted in atherogenesis by the accumulation of intimal immune cells and extracellular lipids. Cap SMCs can be identified by their expression of SMC contractile proteins, such as ACTA2. Lesions also contain large populations of cells that are derived from SMCs but have modulated to other phenotypes, including fibroblast-like and osteochondrogenic-like cells (2,4,5).

The endothelial alignment and layered architecture of cap SMCs suggest that caps may be a *neo-media*, formed by the same mechanisms that drive recruitment and differentiation of medial SMCs in the embryo. Developmental studies in the mouse aorta indicate that embryonic medial SMC differentiation involves an endothelium-elicited Notch signaling cascade that represses the default osteochondrogenic program of the local mesenchymal precursors and steers them towards a SMC fate (6,7).

The Notch family consists of 4 receptors and 5 ligands that provide cell-to-cell communication to coordinate cell fates and spatial arrangements of tissues. Ligand binding leads to cleavage of the intracellular receptor domain (NICD), which translocates to the nucleus where it forms a complex with RBPJ and other factors to drive target gene expression (8).

In the present study, we used conditional knockout of RBPJ and overexpression of NICD to block and force Notch signaling, respectively, in SMCs. We found that Notch signaling is dispensable for the accumulation of SMC-derived cells in plaques, but is required for the re-acquisition of SMC identity in the cap, which forms partly from alternative sources when Notch signaling in medial SMCs is blocked. Conversely, forced Notch signaling inhibited the ability of medial SMCs to contribute to plaque cells and inhibited lesion development. The combined evidence indicates that medial SMCs on their course to become cap SMCs pass through an intermediate cell phenotype with low Notch signaling activity followed by re-acquisition of cap SMC phenotype in a Notch-dependent process that bears similarities to arterial media development.

## Methods

### Mouse experiments

Animal experiments were approved by the ethical review boards at CNIC and Universidad Autónoma, and permitted by the Comunidad de Madrid (PROEX 266/16). Myh11*-*CreER^T2^ mice (B6.FVB-Tg[Myh11-cre/ERT2]1Soff/J, Jackson Laboratory), with tamoxifen-inducible CRE under the SMC-specific *Myh11* promoter (9), were crossed with several mouse strains to generate the following models: floxed TdTomato reporter mice (B6.Cg-*Gt(ROSA)26Sor^tm14(CAG-tdTomato)Hze^*, Jackson Laboratory), containing a loxP-flanked STOP cassette that blocks the transcription of a red fluorescent protein variant (TdTomato) driven by a CAG promoter and inserted in the *GT(ROSA)26Sor* locus (10); *Rbpj^fl/fl^* mice (*Rbpj^tm1Hon^*), constructed with loxP sites flanking exon 6 and 7, which include the essential DNA binding domain (11); and NICD transgenic mice (*Gt(ROSA)26Sor^tm1(Notch1)Dam^*), containing a floxed STOP sequence (LSL) followed by a fragment of the *Notch1* gene encoding the intracellular domain (amino acids 1749-2293, lacking the PEST domain located in the c-terminal region), together with a GFP targeted to the nucleus by an IRES sequence (12). Mouse strains were all backcrossed to the B6 background; SNP matching to the B6 genetic background was >95% for breeders used in the production of experimental groups (Charles River Genetic Testing Service).

All compared mice were littermates, housed together, subjected to the same procedures, and differed only in genotype. All mice included in experiments were males, since Myh11*-*CreER^T2^ is inserted in Y chromosome. In studies of forced Notch signaling, mice hemi- and homozygous for the floxed NICD transgene were grouped together. Recombination was induced in TdTomato reporter mice by two 5-day injection series with 1 mg tamoxifen (T5648-1G, Sigma-Aldrich) per day and in other mice by one 5-day injection series with 2 mg tamoxifen per day (which reported higher recombination efficiencies in our studies).

Hypercholesterolemia was induced 2-4 weeks after the final tamoxifen injection by tail vein injection of rAAV8-mD377Y-mPCSK9 virus particles (1 × 10^11^ vector genomes, produced in the CNIC viral vector core facility) followed by feeding with a high-fat diet (S9167-E011, Sniff, or TD.88137, ENVIGO) (13). Plasma total cholesterol was estimated in duplicate with an enzymatic cholesterol reagent (CH201, Randox Reagents).

Mice were euthanized after 20 weeks of atherosclerosis development by i.p. injection of pentobarbital (250 mg/kg) and lidocain (20 mg/kg) followed by exsanguination and perfusion through the left ventricle with KCl (50 mM, 30 seconds) and 4% phosphate-buffered formaldehyde (5 minutes) at approximately 100 mmHg. Mice were then immersion-fixed in 4% phosphate-buffered formaldehyde for another 18 hours, followed by storage in PBS at 4 °C before vessels were extracted, cryoprotected in sucrose, and frozen in OCT for cryosectioning.

To preserve the GFP signal, mice with the NICD-IRES-GFP transgene were perfusion-fixed for 10 min and not immersion-fixed.

### Tissue processing and staining

Cross-sections (10 μm) of the aortic root were obtained starting from the commissures of the aortic valves. The brachiocephalic trunk was sectioned (10 μm) from the aortic end until no plaques were present. General morphology was assessed by staining sections with Mallory’s trichrome, and specific proteins were detected by immunostaining with the following antibodies: Cy3-coupled mouse anti-ACTA2 (C6198-100UL, 1:500, Sigma-Aldrich) or biotinylated mouse anti-ACTA2 (MS-113-B0, 1:100, Thermo Fisher Scientific) followed by Cy5-streptavidin (SA1011, 1:500, Thermo Fisher Scientific). Rabbit anti-Jag1 monoclonal antibody (2620S, 1:50, Cell Signaling Technology), rabbit anti-Notch3 polyclonal antibody (ab23426, 1:100, Abcam), and rabbit anti-SOX9 (ab185230, dilution 1:100, Abcam) revealed with Alexa Flour 647 Goat anti-rabbit IgG (H+L) (A-21245, 1:500, Thermo Fisher Scientific). Because of the frailty of the GFP signal in NICD-GFP expressing mice, sections from these mice were post-stained for SOX9 after microscopy detection of GFP in ACTA2+ stained sections. Specificity was tested with negative control stains omitting the primary antibody.

### Microscopy and image analysis

All immunostainings were analyzed with a Leica TCS SP5 microscope fitted with a ×40/1.3 Oil objective, except for SOX9 stainings that were analyzed in a Zeiss LSM700 confocal microscope with a 25x/0.8 multi-immersion (corrected for oil) objective. Leica LAS X and ZEN black 2011 software were used for image acquisition, respectively. ImageJ (NIH) was used for quantification. The entire aortic root was scanned by acquiring multiple Z-stack images and stitching them. Plaques in the left coronary sinus were quantified because this region shows more advanced disease in mouse models of atherosclerosis. Nuclei with surrounding ACTA2 signal were counted in the arterial media, plaque, and the cap region, which was defined as the area within 30 μm of the lumen, as described previously (14). Nuclei with SOX9 signal were identified in plaques using an ImageJ macro and checked manually for correctness. TdTomato expression was determined for each of the counted cells in *Rpbj*^SMC-KO^ and *Rpbj*^WT^ mice, whereas GFP expression was determined in NICD^SMC-TG^ mice. The intensity of nuclear GFP expression in cells of mice with the NICD transgene was weak compared with elastin-fiber autofluorescence. To aid analysis, we therefore used the DAPI channel to identify and mask the non-nuclear green channel signal. All acquisition and postprocessing settings were tested with sections from *Myh11-CreER^T2^* controls to confirm specificity of endogenous fluorescence detection. Mallory’s trichrome stained sections were scanned on a digital slide scanner, and plaque area was analyzed with Qupath v.2 software in each aortic root sinus in 3 serial sections (0μm, 100μm, and 200μm from the aortic valve commissures) and presented as the mean area across sections.

### Statistics

Bars in dot blots represent mean ± SEM. P values were calculated by unpaired 2-tailed Student t test for normally distributed data and by 2-tailed Mann-Whitney test for data with a non-normal distribution. Plasma cholesterol burden was quantified as the area under the curve over the course of experiments. Differences in cholesterol level were adjusted for by multiple regression using genotype as a binary variable. The presence of caps was compared between groups by two-tailed chi-square test. Differences were considered statistically significant at *P*<0.05. All statistical tests were performed in Prism 8 (GraphPad Software).

Please see the Supplementary Material for descriptions of additional methods.

## Results

### Notch elements are expressed in the fibrous cap region

During embryonic development of the arterial media, endothelial Jag1 initiates a lateral induction process in which neighboring Notch receptor-expressing mesenchymal cells are induced to differentiate to SMCs. The recruited SMCs are also induced to express Jag1, leading to consecutive rounds of SMC recruitment (7,15).

To analyze whether Notch signaling components could drive fibrous cap formation in atherosclerosis via a similar mechanism, we mapped Jag1 and Notch3, a highly expressed Notch receptor in vascular SMCs (8), in mouse plaques. To unequivocally identify all SMC-derived plaque cells, we used mice carrying a TdTomato Cre reporter transgene and inducible Cre recombinase under the control of the SMC-specific myosin heavy chain promoter (Myh11-CreER^T2^-TdTomato mice). SMC labeling was activated by tamoxifen administration to 6–8-week-old mice, and atherosclerosis was subsequently induced by injecting rAAV-mPCSK9^D377Y^ to reduce hepatic LDL receptor levels and placing the mice on a high-fat diet. After 20 weeks of atherogenesis, Notch3 was expressed in SMCs in the cap region and arterial media, where SMCs also express ACTA2, but was hardly found in modulated SMC-derived cells in the plaque interior (Figure 1). Jag1 was expressed in endothelium and in underlying SMCs and non-SMCs in the cap region, suggesting that cap formation could involve a Notch-mediated lateral induction akin to arterial media development in the embryo. This was supported by in vitro studies showing that increased Notch signaling skewed SMCs to a more contractile phenotype and induced Notch3 and Jag1, consistent with published findings (**Supplementary Figure 1**) (7,16).

**Figure 1.**
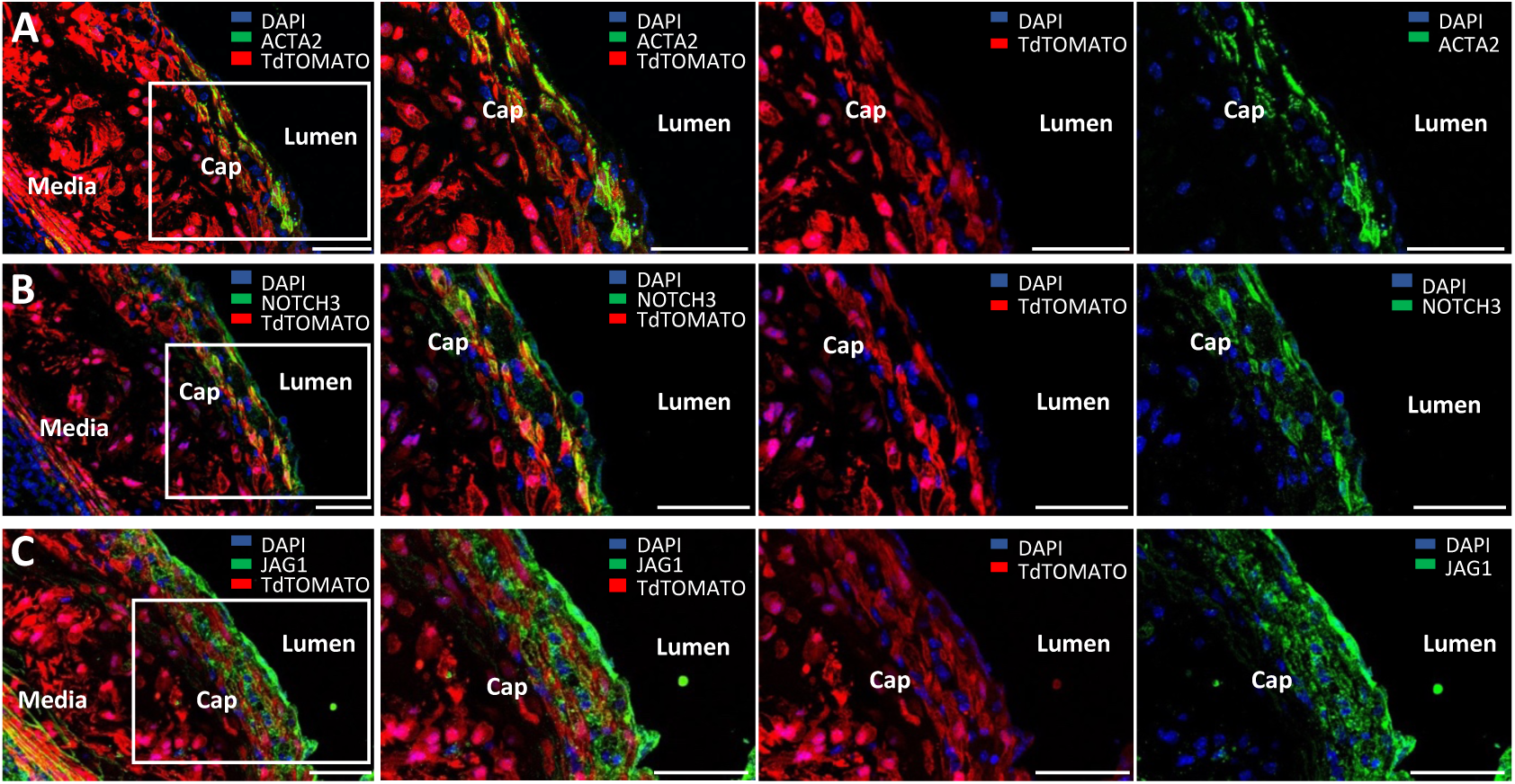
Notch signaling components in areas of fibrous cap formation. (**A**) Representative aortic root section showing atherosclerotic plaque in the left coronary sinus of a Myh11-CreERT2-TdTomato mouse with cap ACTA2+ SMCs under the endothelium and ACTA2-SMC-derived cells (TdTomato+) in the plaque interior. (**B-C**) Consecutive sections showing the expression of Jag1 and Notch3 in the cap region. Scale bars, 50 μm.

### Activation of Notch signaling is required for fibrous cap SMC fate in plaques

To test the importance of Notch signaling for cap formation, we analyzed atherosclerosis development in mice lacking Notch signaling in SMCs due to loss of the essential Notch effector transcription factor RBPJ. We bred littermate Myh11-CreER^T2^-TdTomato mice that had either homozygous floxed *Rbpj* alleles (*Rbpj*^SMC-KO^, n=20) or the wildtype alleles (*Rbpj*^WT^, n=16). Tamoxifen was injected in 6-week-old mice to recombine and inactivate the floxed *Rbpj* alleles, and atherosclerosis was subsequently induced by rAAV-mPCSK9^D377Y^ injection and the 20-week high-fat diet. *Rbpj*^SMC-KO^ mice appeared healthy, and their arteries had a grossly normal appearance (**Supplementary Figure 2**).

Atherosclerosis was analyzed in the aortic root region (Figure 2A-B). SMC-specific disruption of *Rbpj* did not alter plasma cholesterol levels or mean plaque size in the total root. Recruitment of cap SMCs was analyzed by high-resolution microscopy in the left coronary sinus, where the most advanced lesions with fibrous caps form in mice. Plaque size was found to be moderately increased in the left coronary sinus of *Rpbj*^SMC-KO^ mice; however, despite the larger plaques, the number of ACTA2+ cap cells in the subendothelial region (defined as <30 μm from the lumen) was significantly reduced (Figure 2C). *Rpbj*^SMC-KO^ mice usually had thinner, one-layer caps, whereas *Rpbj*^WT^ mice frequently developed thicker caps with several layers of ACTA2+ SMCs (Figure 2D). Moreover, 30% of analyzed *Rpbj*^SMC-KO^ mice had no fibrous caps in the left coronary sinus (less than 2 ACTA2+ cells found together), whereas cap formation was detected in all *Rpbj*^WT^ mice (*P*<0.05, 2-sided chi-square test).

**Figure 2.**
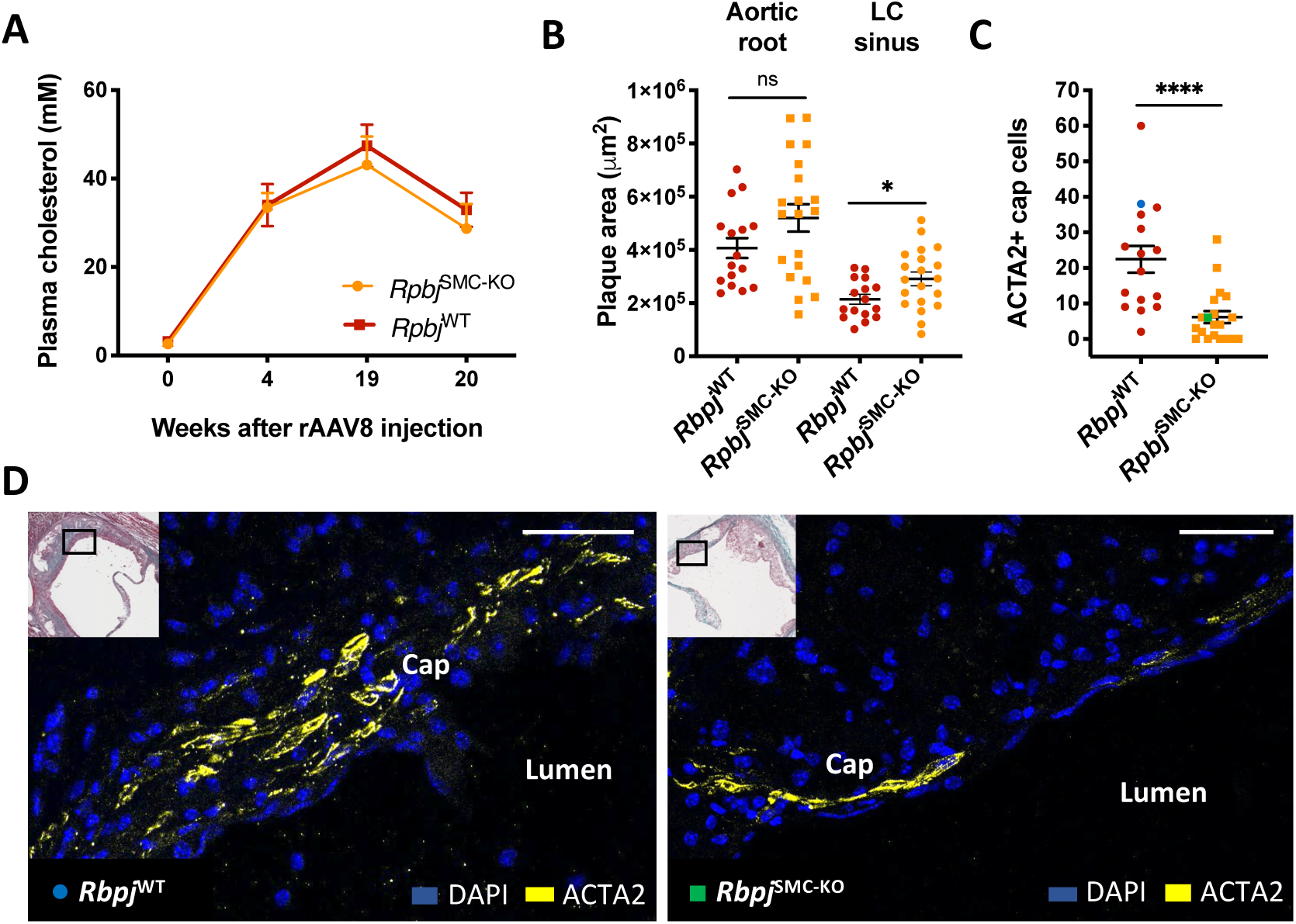
Inactivating Notch signaling in SMCs reduces cap formation. (**A**) Total plasma cholesterol in *Rbpj*^WT^ and *Rbpj*^SMC-KO^ mice after rAAV-mPCSK9^D377Y^ injection and a high-fat diet (bars show mean ± SEM). (**B**) Plaque size in the total aortic root and the atherosclerosis-prone left coronary (LC) sinus (\**P*<0.05, unpaired t-test). (**C**) Number of ACTA2+ cap cells (<30 μm from the endothelium) in *Rbpj*^WT^ and *Rbpj*^SMC-KO^ mice (lacking Notch signaling in SMCs) (\*\*\*\**P*<0.0001, Mann-Whitney test). Blue and green dots correspond to the examples shown in E and F. (**D**) Representative ACTA2+ staining in left coronary sinus sections from *Rbpj*^WT^ and *Rbpj*^SMC-KO^ mice, showing a less developed, thinner cap in *Rbpj*^SMC-KO^ mice and thicker cap in *Rbpj*^WT^ mice. Scale bars, 50 μm.

The reduction in cap SMCs in *Rpbj*^SMC-KO^ mice could be caused by either reduced recruitment to the cap region or by phenotypic changes in the recruited cells. To distinguish between these possibilities, we counted all SMC-derived plaque cells in *Rpbj*^SMC-KO^ and *Rpbj*^WT^ mice using the TdTomato lineage tracer. No differences were detected in TdTomato+ cell numbers in the cap region or across the whole plaque (Figure 3A-B). We also found no differences in the recruitment of SMC-derived SOX9-expressing osteochondrogenic plaque cells (Figure 3C-E). These data indicate that Notch signaling is dispensable for the accumulation of SMC-derived cells in plaques, but is required for SMC phenotype acquisition in the cap. This conclusion was strongly supported by the observation that many of the small number of ACTA2+ cap SMCs in *Rpbj*^SMC-KO^ mice were recruited from cells that had not recombined the easily recombinable TdT reporter, and therefore by inference would also not have recombined the more difficult floxed *Rbpj* gene (Figure 3F-I). These results thus indicate that cells with intact Notch signaling have an important competitive advantage in the formation of ACTA2+ cap SMCs.

**Figure 3.**
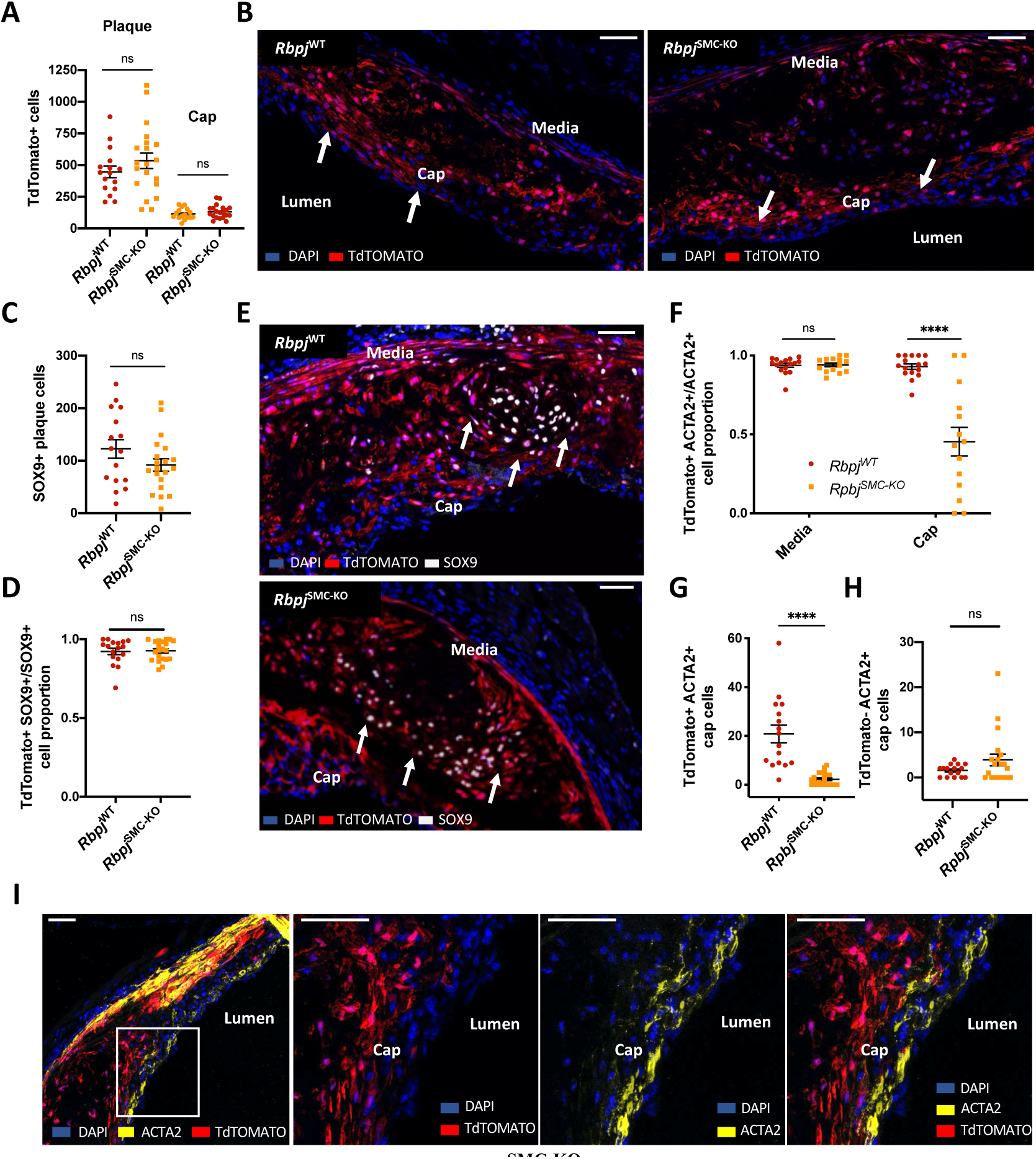
SMC-derived plaque cells in *Rbpj*SMC-KO mice accumulate in the cap region but fail to acquire SMC phenotype. (**A-B**) Number of SMC-derived (TdTomato+) cells in the full plaque and specifically in the cap region in *Rbpj*^WT^ and *Rbpj*^SMC-KO^ mice. Arrows in B point to TdTomato+ cells in the cap region. (**C-E**) Recruitment of SOX9+ osteochondrogenic plaque cells from recombined (TdTomato+) medial SMCs. Panels show total SOX9+ plaque cells (C), the proportion of these cells positive for the SMC lineage tracer (D), and representative stainings (E). Arrows point to SOX9+ cells. (**F**) Contribution of recombined medial SMCs to cap SMCs in *Rbpj*^WT^ and *Rbpj*^SMC-KO^ mice. Most medial SMCs expressed TdTomato in both groups, but significant differences were found in the cap, where ACTA2+ cap cells in *Rbpj*^SMC-KO^ mice were frequently TdTomato-(\*\*\*\**P*<0.0001, unpaired t-test). (**G-H**) Cap TdTomato+ ACTA2+ cells (G) and TdTomato– ACTA2+ cells (H) in *Rbpj*^WT^ and *Rbpj*^SMC-KO^ mice, showing a significant reduction in recruitment of TdTomato+ cells in *Rbpj*^SMC-KO^ mice (\*\*\*\**P*<0.0001, Mann-Whitney test) and a numerical, albeit non-significant, increase in the number of cap ACTA2+ SMCs recruited from nonrecombined (TdTomato–) cells. (**I**) Example of cap formation in an *Rbpj*^SMC-KO^ mouse in which most ACTA2+ cap SMCs were recruited from cells that had not recombined the TdTomato reporter.

Additional analysis was performed in the brachiocephalic trunk. The development of atherosclerosis at this site was much more variable, and no significant differences were detected in the total number of ACTA2+ cells. However, again significantly more of the ACTA2+ cells were recruited from TdT-negative cells in *Rpbj*^SMC-KO^ mice than in *Rbpj*^WT^ mice (**Supplementary Figure 3**).

### Down-regulation of Notch signaling is necessary for accumulation of SMC-derived plaque cells

Unaltered recruitment of SMC-derived plaque cells in *Rpbj*^SMC-KO^ mice does not necessarily imply that Notch signaling is unimportant for the contribution of SMCs to plaque cells. This finding could also indicate that the Notch pathway is already shut down in SMCs that are recruited and undergo clonal expansion in plaque. In such a scenario, knockout of *Rbpj* would be inconsequential to the accumulation of SMC-derived plaque cells.

To test the possibility that Notch is required to be down-regulated for medial SMCs to contribute to plaque and cap formation, we turned to Myh11-CreER^T2^ LSL-NICD-GFP mice (NICD^SMC-TG^). SMCs in these mice can be induced to constitutively express Notch intracellular domain, thereby abolishing their ability to downregulate Notch signaling. Recombination and atherosclerosis induction were performed in NICD^SMC-TG^ (n=17) and littermate Myh11-CreER^T2^ control mice (n=14) as described above for *Rbpj*^WT^ and *Rbpj*^SMC-KO^ mice.

Constitutive Notch signaling in SMCs substantially reduced the development of atherosclerotic lesions in the total aortic root and specifically in the atherosclerosis-prone left coronary sinus (Figure 4A-B). Unexpectedly, plasma cholesterol levels ended up being moderately lower in NICD^SMC-TG^ mice than in controls, but this difference was insufficient to explain the differences in plaque development (Figure 4C-D), which remained significant after correcting for the differences in cholesterol burden in a multiple regression model (**Supplementary Table 1**).

**Figure 4.**
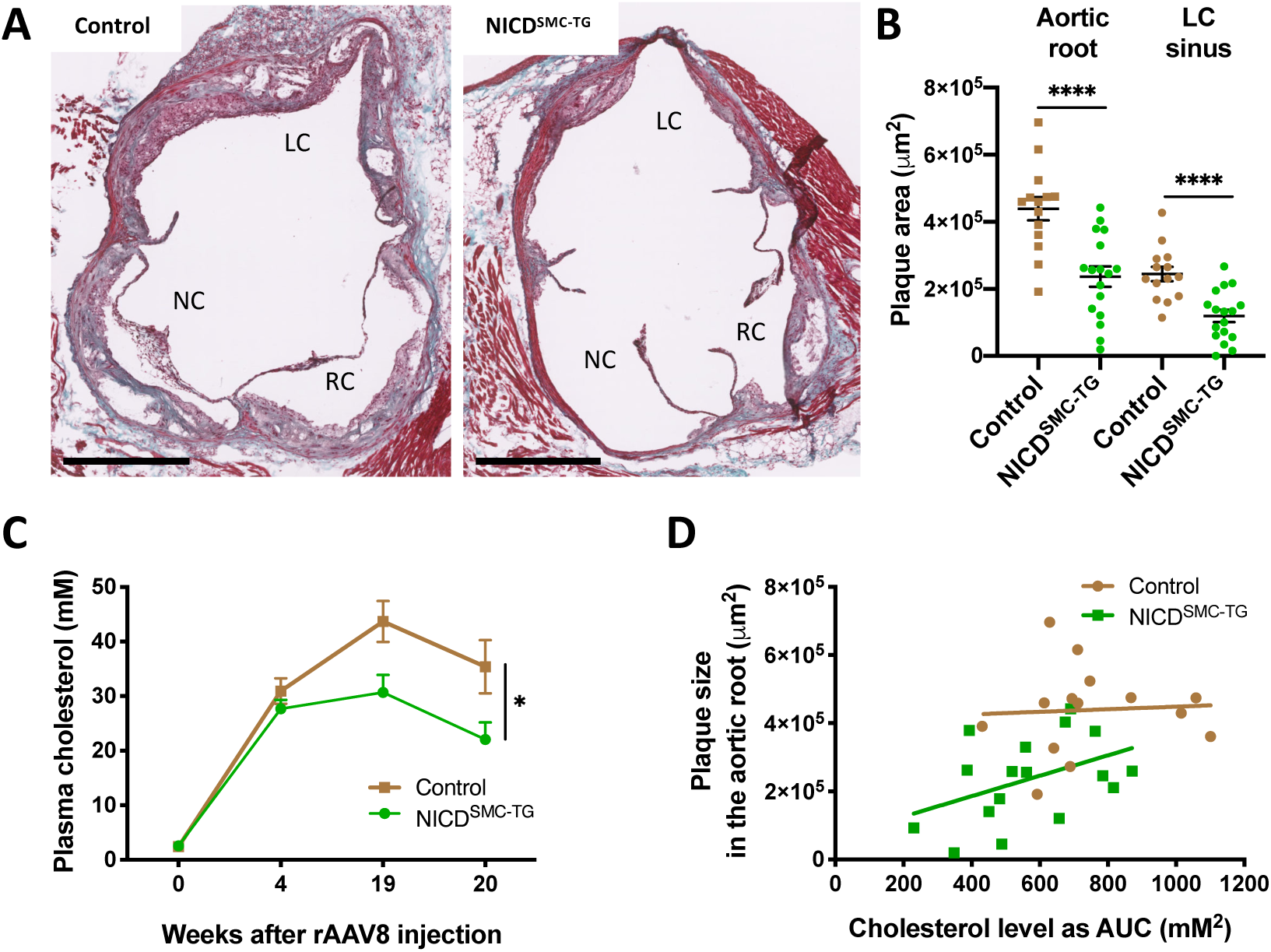
Constitutive Notch activation in SMCs reduces plaque formation. (**A**) Representative aortic root sections showing plaques in NICD^SMC-TG^ and control mice. Mallory’s trichrome staining. Scale bars, 500 μm. LC, Left coronary sinus; RC, Right coronary Sinus; NC, Non-coronary sinus. (**B**) Plaque size measured in the total aortic root and specifically in the left coronary (LC) sinus (\*\*\*\**P*<0.0001, unpaired t-test). (**C**) Total plasma cholesterol in NICD^SMC-TG^ and control mice after rAAV-mPCSK9^D377Y^ injection and a high-fat diet (bars show mean ± SEM). Note the similar profiles up to week 4, after which NICD^SMC-TG^ mice had significantly lower plasma cholesterol than their littermate controls (\**P*<0.05, unpaired t-test of area-under-the-curve) (**D**) Relationship between plaque size and plasma cholesterol burden measured as area under the curve in NICD^SMC-TG^ and control mice. The upward displacement of the regression lines in the two groups indicates that the difference in plaque size was not explained by the differences in cholesterol levels. The slope of both regression lines was not significantly different from 0.

Despite the differences in plaque size, plaques in NICD^SMC-TG^ mice had normal numbers of total ACTA2+ SMCs and cap ACTA2+ SMCs (Figure 5A). To determine whether these cap SMCs were derived from SMCs with constitutive Notch activation, we contrasted the fate of recombined and non-recombined SMCs in plaques from NICD^SMC-TG^ mice. Recombined SMCs were identified by the expression of nuclear GFP and constituted 60.3±3.2% (mean ± SEM) of medial ACTA2+ SMCs in NICD^SMC-TG^ mice. Nevertheless, the proportion of GFP+ ACTA2+ cells in the plaque was significantly lower, at 14.1±2.4% (Figure 5B-C), indicating that forced Notch signaling inhibited the ability of SMCs to contribute to cap formation. A similar analysis of osteochrondogenic SOX9+ plaque cells, which are also recruited from medial SMCs, revealed almost no contribution from GFP+ NICD-expressing cells, at 2.7±0.9% (Figure 5B, D). Down-regulation of Notch signaling, which is impossible in NICD-expressing cells, thus appear to be mandatory for medial SMCs to generate SMC-derived cell populations in atherosclerotic plaques.

**Figure 5.**
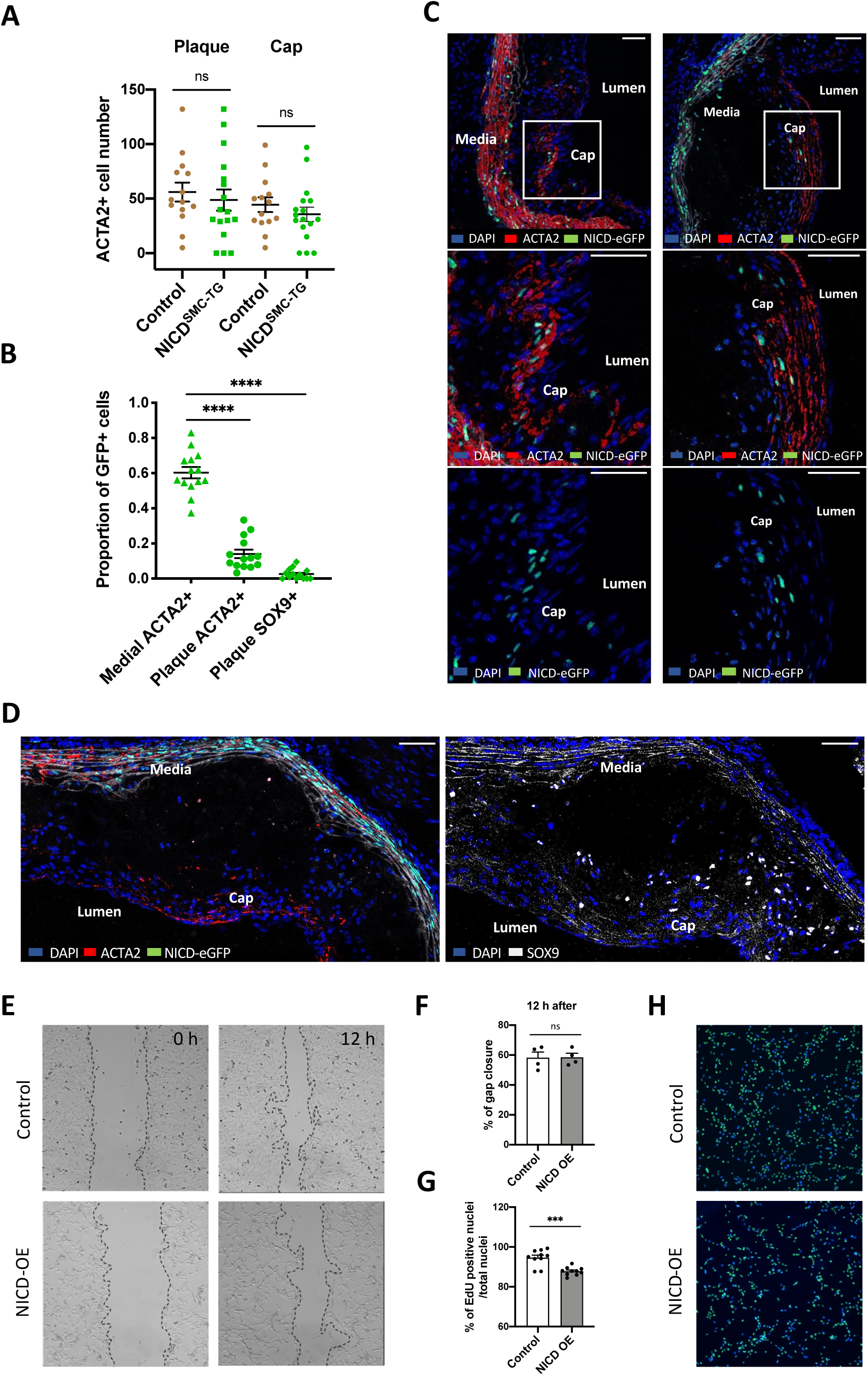
Notch signaling activation in SMCs limits their contribution to atherogenesis. **(A)** Number of ACTA2+ SMCs in the plaque and the cap region (<30 µm from the endothelium) in NICD^SMC-TG^ and control mice, showing no between-genotype differences despite the difference in plaque size. (**B**) Fraction of GFP+ cells among ACTA2+ medial and plaque cells and SOX9+ plaque cells. The significantly lower proportion of GFP+ cells among plaque SMCs and osteochondrogenic cells than among medial SMCs in NICD^SMC-TG^ mice indicates that NICD overexpression inhibited the ability of medial SMCs to contribute to plaque ACTA2+ cells and, even more so, to osteochondrogenic SOX9+ cells (\*\*\*\**P*<0.0001, unpaired t-test). (**C**) Two representative sections showing ACTA2+ staining in plaques from NICD^SMC-TG^ mice. A high proportion of ACTA2+ cells in the media are GFP+, but in the plaque only a few recruited ACTA2+ cells are GFP+ (and therefore overexpressing NICD). (**D**) Representative immunostaining for ACTA2 (left, performed first) and SOX9 (right, performed after microscopy acquisition) on an NICD^SMC-TG^ aortic root section; GFP+ ACTA2+ cells are frequent in the media, but contribute little to the SOX9+ osteochondrogenic population in the plaque. (**E-F**) In vitro SMC migration assay. NICD overexpression (NICD-OE) in rat SMCs did not alter migration capacity in scratch assays, but cells were less dense and more spindle-shaped. (**G-H**) Proliferation of rat SMCs assessed by EdU incorporation, showing significantly reduced proliferation in NICD-OE SMCs (\*\*\**P*<0.01, unpaired t-test). Scale bars, 50 µm.

Accumulation of SMC-derived cells in plaque involves migration of a small number of medial SMCs and their subsequent proliferation (2,17). To explore which of these steps is inhibited by constitutive Notch signaling, we electroporated cultured rat SMCs with an NICD expression plasmid and assayed them for migration and proliferation. Constitutive Notch signaling did not significantly inhibit migration in scratch assays (Figure 5E-F); however, cells were more dispersed, spindle-shaped, and they proliferated less than cells electroporated with empty vector (Figure 5G-H).

Interestingly, the impaired proliferation of NICD-expressing SMCs in vitro was consistent with the lack of large GFP+ clones in NICD^SMC-TG^ plaques. Previous studies have shown that SMC-derived cells accumulate in plaque by clonal expansion of a few medial SMC precursors (2,17). Thus, if NICD overexpression did not influence clonal expansion, we would expect some of the recruited GFP+ cells to give rise to large GFP+ clones, but this was not observed. Only small GFP+ clusters could be seen (examples in Figure 5C), and most GFP+ cells were dispersed in the plaque, consistent with an inability of NICD-expressing SMCs to undergo clonal proliferation.

## Discussion

In the present study, we show that Notch signaling is a key regulator of SMCs during atherosclerosis and fibrous cap formation. Notch signaling is required to be downregulated for medial cells to mobilize and expand in lesions. Conversely, it is required to be subsequently activated for some of these accumulating cells to re-acquire SMC phenotype in the cap.

The involvement of Notch signaling in driving cells towards a SMC phenotype in the subendothelial space bears strong resemblance to arterial media development. During the development of the aortic arch, medial SMCs are formed when Jag1 expressed on endothelial cells activates the Notch pathway in neighboring neural crest mesenchymal cells (7,15). The first recruited SMCs themselves express Jag1 and can thus recruit the next layer of medial SMCs, thereby providing a mechanism for building the multilayered arterial media (7). Notch signaling also plays a decisive role in the embryonic distal aorta, where Notch signaling is the critical factor that switches mesodermal mesenchymal cells away from a SOX9-driven osteochondrogenic fate and towards SMC differentiation. The resemblance between these embryonic processes and cap formation is underscored by the phenotypic spectrum of SMC-derived cells in atherosclerosis; recent single-cell expression profiling of SMC-derived cells in atherosclerosis and related pathologies indicates that they broadly recapitulate the cell types derived from embryonic mesenchyme, including SMCs, fibroblast-like cells, and osteochondrogenic-like SOX9+ cells (4,18). Together, these observations suggest that the cap might form from dedifferentiated SMCs as a *neo-media* to substitute the lost connection between the endothelium and the arterial media at sites of plaque development. Creation of this neo-media serves to stabilize the arterial lining against rupture in regions of plaque, as the media serves to stabilize newly developed arteries in development.

Our conclusions are based on inducible SMC conditional knockout and overexpression experiments in mice. There are a number of factors that must be considered when interpreting this type of experiment. First, SMC recombination is typically incomplete. Even the very easily recombined TdTomato transgene (19) was not induced in all medial SMCs, and in the NICD overexpression experiment, analysis of the co-expressed nuclear GFP signal indicated an average recombination rate of approximately 60%. Second, the whole population of SMC-derived cells in plaques is recruited from a low number of medial SMCs. Our previous study in mice with PCSK9-induced atherosclerosis indicated that only 10-12 medial SMCs contribute to the entire SMC-derived cell population in aortic root plaques (2). If there is selective pressure against the recombined medial SMCs, there is ample chance that this small number of cells could be recruited from among the residual non-recombined cells, therefore leading to normal plaque development. In this scenario, contrasting the behavior of recombined and non-recombined cells in the same plaques is the most powerful approach. This provides a direct comparison of the cell-autonomous function of Notch signaling among cells placed in the same milieu.

In *Rpbj*^SMC-KO^ mice, we found that TdT-negative cells were at an advantage in contributing to ACTA2+ cap SMCs but not to osteochondrogenic plaque cells. These results show that when Notch signaling in most medial SMCs is blocked, the cap forms partly from alternative sources with low Myh11-CreER^T2^ activity that are very unlikely to have recombined their *Rbpj* alleles. This alternative source of cap SMCs could be medial SMCs with low *Myh11* promoter activity. It could also be non-SMCs, such as adventitial Sca1+PDGFR*α*+ cells, which have been shown to contribute to neointimal SMCs in situations where medial SMCs are incapacitated (20), or endothelial cells through endothelial-mesenchymal transition (21). Tracking these sources in *Rbpj*^SMC-KO^ mice is challenging because of the need to use lineage tracing techniques that work independently of the Cre recombinase used to recombine the *Rbpj* allele.

Recruitment of ACTA2+ SMCs in the plaque is known to be regulated by several other SMC genes, including *Klf4* (5) and *Oct4* (22). The specific roles of these genes in building the cap are, however, not easy to decipher because both genes exert strong effects on plaque formation and in the accumulation of SMC-derived cells in plaques (5,22,23). A particular strength of the *Rbpj*^SMC-KO^ experiment was that overall recruitment and expansion of SMC-derived plaque cells was unaffected, or affected only to a minor degree. This allowed us to identify the importance of Notch signaling for the generation of ACTA2+ cap SMCs free of biases incurred by differences in the recruitment of SMC-derived cells to the cap region.

### Limitations

Our study does not consider which specific Notch ligands and receptors orchestrate cap formation. Jag1 and Notch3 are strong candidates due to their expression in plaque endothelium and cap SMCs. However, other ligands and receptors may be involved, and both Notch2 and Notch3 receptors are expressed in fibrous cap regions of human plaques (24). To avoid problems caused by potential redundancy of different Notch receptors in SMCs, we carried out experiments with *Rbpj* mutants to examine the full requirement of Notch for cap formation irrespectively of the upstream Notch receptor.

### Perspective

Rupture of thin fibrous caps is the final event that triggers approximately 70% of heart attacks precipitated by coronary atherosclerosis (1). Why some lesions fail to develop or maintain thick caps is not yet understood. If cap formation is analogous to arterial media development, the mechanisms determining media thickness in development may also determine plaque thickness in the adult. Unfortunately, these mechanisms are not yet well understood. Arterial media thickness is tightly controlled to match arterial tensile stress (25), but it is not yet understood what mechano-transduction mechanism regulates the termination of Notch-mediated lateral induction of SMC differentiation and the transition to adventitial fibroblast differentiation (15). Exploring these mechanisms in embryological development and their potential perturbation by the inflammatory and metabolic processes in the atherosclerotic lesion may provide future clues to the cause of thin-cap fibroatheroma formation.

### Conclusion

Sequential loss and gain of Notch signaling is necessary for building the contractile cap SMC population. The similarity of mechanisms with embryonic arterial media assembly suggests that the fibrous cap may form as a *neo-media* to reestablish the connection between endothelium and SMCs in the developing plaque.

## Acknowledgments

We thank the CNIC Viral Vectors Unit, Microscopy Unit, and Animal Facility at CNIC for excellent technical help, and Simon Bartlett, CNIC, for English editing.

