## Supplementary material for "Fibrous caps in atherosclerosis form by Notch-dependent mechanisms common to arterial media development"

### Supplementary methods

#### *Rat aortic smooth muscle cell culture*

Rat aortic smooth muscle cells (ATCC CRL-2018) were cultured in DMEM supplemented with 10% FBS, 1% penicillin/streptomycin, and 2 mM glutamine in a humidified incubator at 37 °C with 5% $CO_2$. Cells were used at passages 4-8.

#### *Rbpj siRNA-mediated knock-down*

Rat aortic smooth muscle cells at 70% confluence were transfected with 10 µM control siRNA (AllStars Negative siRNA, Qiagen 1027280) and 10 µM Rbpj siRNAs using Opti-MEM (Thermo Fisher) and Lipofectamine RNAiMAX (Invitrogen) for 18 h on two consecutive days. The rat siRNAs targeting Rbpj as follows: Rn_LOC679028_1 FlexiTube siRNA and Rn_LOC679028_2 FlexiTube siRNA (Qiagen).

#### *NICD overexpression in rat aortic smooth muscle cells*

NICD cDNA was cloned into a mammalian gene expression vector (pRP-CAG>HA/Notch1_NICD; VB200116-1403yad; Vectorbuilder). Rat aortic smooth muscle cells (100,000 cells) were transfected by electroporation (Neon, Thermo Fisher Scientific) with either 1 µg NICD vector (5743 bp) or pcDNA3.1 (empty vector). The electroporation parameters were 1250 V, 20 msec, and 2 pulses. After transfection, cells were cultured in a 24-well plate containing medium without antibiotics. After 24 h, cells were allowed to recover in full medium and were then used in scratch assays (migration), labeled with EdU (proliferation), or harvested (gene expression).

#### *Proliferation assay*

Rat aortic SMCs were seeded at a concentration of 100,000 cells/cm$^2$ on coverslips coated with collagen type I on a 24-well plate. After transfection, cells were then grown for 24 h in medium containing 10 µM EdU. Cells were then fixed for 15 min in 3.7% paraformaldehyde in phosphate-buffered saline (PBS), pH 7.4. EdU-positive cells were then labeled according to the manufacturer's instructions (BCK-EDU488, Sigma-Aldrich), followed by labeling of nuclei with Dapi. A total of two-three independent experiments were performed, each with three-five technical replicates. Stained coverslips were analyzed under an Eclipse Ti2 inverted microscope (Nikon) fitted with a 10X objective. Nikon NIS-Elements software was used for image acquisition, and ImageJ (NIH) was used for quantification.

***Scratch assay***

SMCs were seeded in 24-well plates. The cell monolayer was scratched using a 10 µl pipette tip before washing three times with PBS to clear cell debris and floating cells. The cell plate was mounted under an Eclipse Ti2 inverted microscope (Nikon) and incubated in an integrated chamber for 24 h at 37 °C in 5% $CO_2$. Time-lapse images of the same field were acquired every hour for 24 h. Migration was measured by calculating the percentage of the denuded area covered by cells after 24 h relative to the area at time 0 using NIS-Elements software (Nikon). Two independent experiments were performed, each with two or three technical replicates.

***Quantitative real-time PCR analysis***

Total RNA was isolated from rat smooth muscle cells using the Nucleospin RNA plus kit (Macherey Nagel, Düren, Germany). Complementary DNA synthesis was performed using 0.5 µg RNA with the RevertAid First Strand cDNA synthesis kit (Thermo Scientific). For Quantitative real-time PCR, 100 ng cDNA served as the template for PCR amplification using the Maxima SYBR® Green QPCR Master Mix (Thermo Scientific) in an Aria Mx3000P qPCR System (Agilent Technologies, Santa Clara, CA), in order to detect expression levels of several genes involved in the contractile phenotype and Notch signaling pathway (**Supplementary Table 2**). Three independent experiments were performed, each with three technical replicates. Each reaction was run in duplicate, and relative gene expression levels were normalized to rat hypoxanthine-guanine phosphoribosyltransferase (Hprt1). Relative expression was calculated using the ΔΔCt method.

***PCR to detect Rbpj recombination***

To detect the recombined allele in *Rbpj*$^{SMC\text{-}KO}$ mice, we performed PCR on mouse aorta DNA to amplify either the floxed allele (DY223/DY225, 430bp) or the recombined allele (DY223/DY229, 640bp). The primer sequences were as follows: DY223 (forward) 5’-ACC AGA ATC TGT TTG TTA TTT GCA TTA CTG-3’; DY225 (reverse) 5’-ATG TAC ATT TTG TAC TCA CAG AGA TGG ATG-3’; DY229 (reverse) 5’-TAA TGC ACA CAA GCA TTG TCT GAG TTC-3’.

**Supplementary Figure 1**

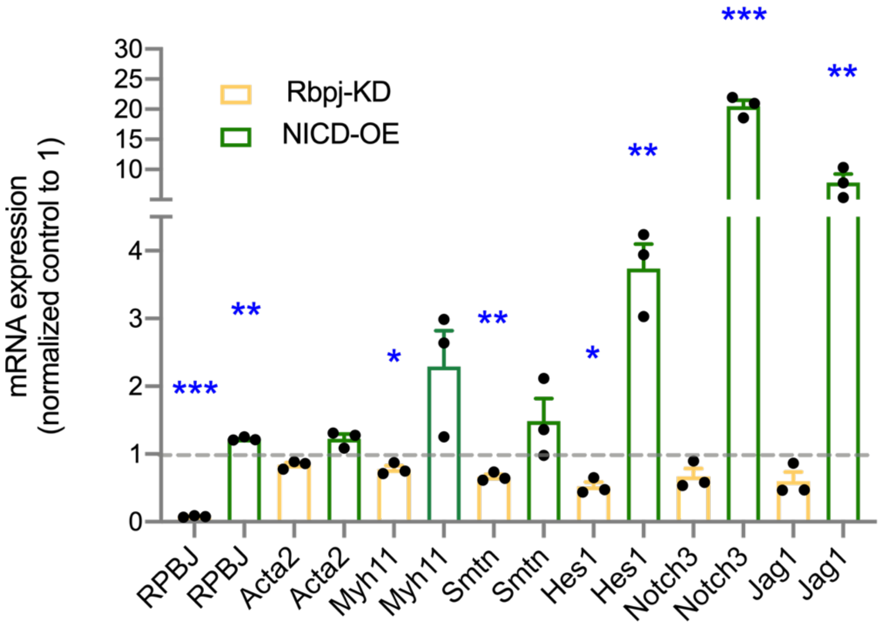

**Supplementary Figure 1. Notch signaling and SMC gene expression.** Notch signaling blockade by *Rbpj* knockdown (Rbpj-KD) and forced Notch signaling by overexpression of a Notch intracellular domain (NICD-OE) regulate contractile SMC genes (Acta2, Myh11, Smtn), Notch targets (Hes1), and Notch elements (Notch3, Jag1) in rat SMCs. Each point represents the mean expression relative to controls in an independent experiment. *p<0.05, **p<0.01, ***p<0.001, unpaired t-tests.

**Supplementary Figure 2**

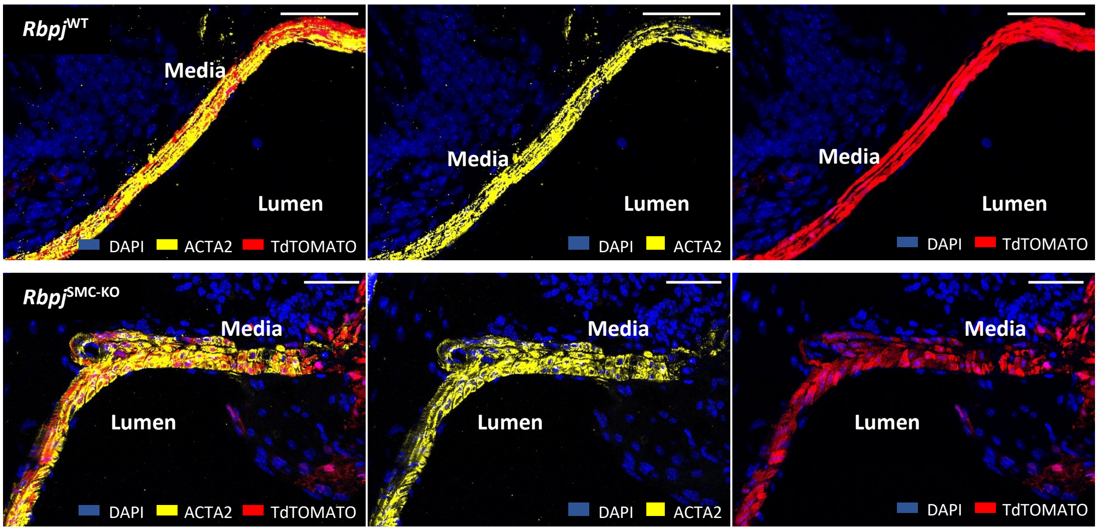

**Supplementary Figure 2. Examples of arterial morphology in *Rbpj*$^{SMC\text{-}KO}$ *and Rbpj*$^{WT}$ mice.** In arteries not affected by atherosclerosis, the contractile phenotype of medial SMCs was conserved as indicated by positive staining for ACTA2. Examples are from the beginning of the left coronary artery. Scale bars, 50 µm.

### Supplementary Figure 3

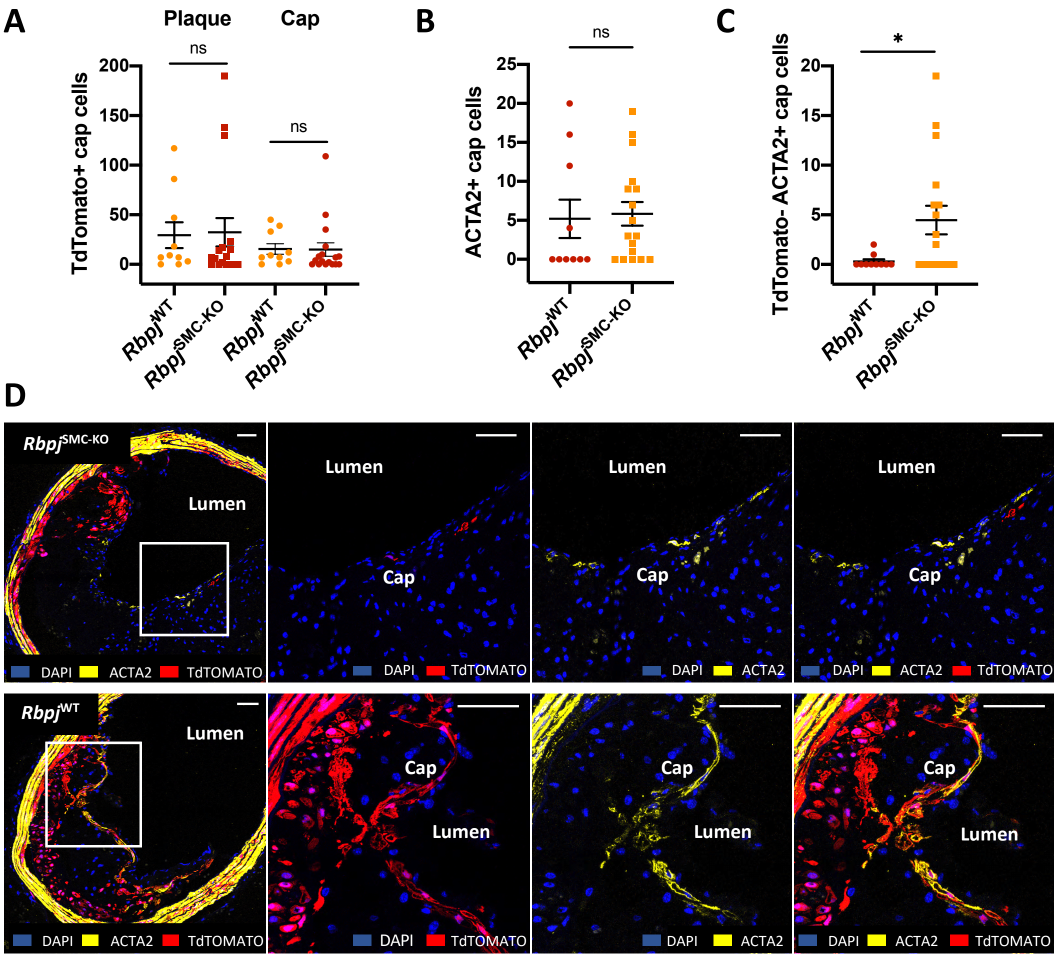

**Supplementary Figure 3. Notch inactivation in SMCs impairs their ability to contribute to cap in the brachiocephalic trunk.** (**A**) The number of TdTomato+ cells in plaque and specifically in the cap region (<30 µm from the endothelium) did not differ between *Rbpj*$^{SMC\text{-}KO}$ and *Rbpj*$^{WT}$ mice. (**B**) Number of ACTA2+ cells in the cap region was also similar. (**C**) Analysis of whether these ACTA2+ cells were recruited from medial SMCs that had recombined the TdTomato reporter, however, revealed major differences. In *Rbpj*$^{SMC\text{-}KO}$ mice, significantly more ACTA2+ were recruited from TdTomato- cells, indicating a competitive advantage of cells that had intact Notch signaling compared with cells that had not. (**D**) Representative examples of ACTA2 staining of BCT plaque from *Rbpj*$^{SMC\text{-}KO}$ and *Rbpj*$^{WT}$ mice. The plaque from the *Rbpj*$^{SMC\text{-}KO}$ mouse shows ACTA2+ TdTomato- cells, whereas ACTA2+ cells are TdTomato+ in plaque from the *Rbpj*$^{WT}$ mouse. Scale bar, 50 µm.

**Supplementary Table 1**

**Regression model**

$Y = \boldsymbol{\beta_1} \cdot \mathrm{B} + \boldsymbol{\beta_2} \cdot \mathrm{C} + \boldsymbol{\beta_0}$

where Y is plaque size in the LC sinus of the aortic root after 20 weeks of atherogenesis (in $\mu m^2$), B is a binary variable representing genotype, and C is the area under the curve of plasma cholesterol levels from week 0 until week 20 (in mM).

| | **Estimate** | **P value** |
|---|---|---|
| $\boldsymbol{\beta_0}$ | 308481 (95% CI: 105117 to 511845) | 0,0043 |
| $\boldsymbol{\beta_1}$ | -170961 (95% CI: -274478 to -67445) | 0,0021 |
| $\boldsymbol{\beta_2}$ | 174,2 (95% CI: -81,04 to 429,5) | 0,1721 |

**Supplementary Table 1. Multiple regression for plaque size determinants in NICD$^{SMC\text{-}TG}$ and control mice.** Only genotype was independently associated with plaque size in a model that also incorporated plasma cholesterol burden. This indicates that the reduced atherogenesis observed in NICD$^{SMC\text{-}TG}$ mice compared with control mice cannot be explained by the moderate difference in plasma cholesterol levels between groups.

**Supplementary Table 2**

| Gene | Forward | Reverse |
|---|---|---|
| Notch3 | 5’-CCTCCTGAAGAGTTGCTTCG-3’ | 5’-CGAGGTGCGCAGAATAGC-3’ |
| Jag1 | 5’-CGCCCTCTGAAAAACAGAAC-3’ | 5’-ACCCAAGCCACTGTTAAGACA-3’ |
| Rbpj | 5’-GAGCGTCCTGGACAATCATC-3’ | 5’-GGCCCATTCCCTCATAGAAC-3’ |
| Hairy and enhancer of split-1 (Hes1) | 5’-AGTGAAGCACCTCCGGAAC-3’ | 5’-CGTTCATGCACTCGCTGA-3’ |
| Smoothelin (Smtn) | 5’-CACATTGGACCCTGGTAAGG-3’ | 5’-GTCCTCTGGTGTCGGAGTTG-3’ |
| Myosin-11 (Myh11) | 5’-TCAATCACACCATGTTCATCC-3’ | 5’-CACCAGGAGGGTTGTTCG-3’ |
| Actin alpha 2 (Acta2) | 5’-CCCAGCACCATGAAGATCA-3’ | 5’-CGCCGATCCAGACAGAAT-3’ |

**Supplementary Table 2. Primer sequences for quantitative real-time PCR analysis.** Forward and reverse primer sequences for the amplification of Notch-related genes and contractility markers.
